# Beyond self-resistance: ABCF ATPase LmrC is a signal-transducing component of an antibiotic-driven signaling cascade hastening the onset of lincomycin biosynthesis

**DOI:** 10.1101/2020.10.16.343517

**Authors:** Marketa Koberska, Ludmila Vesela, Vladimir Vimberg, Jakub Lenart, Jana Vesela, Zdenek Kamenik, Jiri Janata, Gabriela Balikova Novotna

**Affiliations:** Institute of Microbiology v. v. i., The Czech Academy of Sciences, BIOCEV, Vestec, Czech Republic; Charles University in Prague, Faculty of Science, Department of Genetics and Microbiology, Prague, Czech Republic; Institute of Microbiology v. v. i., The Czech Academy of Sciences, Prague, Czech Republic

## Abstract

In natural environments, antibiotics are an important instrument of inter-species competition. At subinhibitory concentrations, they act as cues or signals inducing antibiotic production: however, our knowledge of well-documented antibiotic-based sensing systems is limited. Here, for the soil actinobacterium *Streptomyces lincolnensis* we describe a fundamentally new ribosome-mediated signaling cascade that accelerates the onset of lincomycin production in response to an external ribosome-targeting antibiotic to synchronize the antibiotic production within the population. The entire cascade is encoded within the lincomycin biosynthetic gene cluster (BGC) and besides the transcriptional regulator, LmbU it consists of three lincomycin resistance proteins: a lincomycin transporter, LmrA, a 23S rRNA methyltransferase, LmrB, both conferring a high resistance, and an ABCF ATPase LmrC that confers only moderate resistance but is indispensable for the antibiotic-induced signal transduction. Specifically, the antibiotic sensing occurs via a ribosome-mediated attenuation, which activates LmrC production in response to lincosamide, streptogramin A, or pleuromutilin antibiotics. Then, the ribosome-operating LmrC ATPase activity triggers the transcription of *lmbU* and consequently the expression of lincomycin BGC. Finally, the production of LmrC is downregulated by LmrA and LmrB which reduces the amount of the ribosome-bound antibiotic and thus fine-tune the cascade. We propose that analogous ABCF-mediated signaling systems are relatively common because many BGCs for ribosome-targeting antibiotics encode an ABCF-protein accompanied by additional resistance protein(s) and transcriptional regulators. Moreover, we revealed that three of eight co-produced ABCF proteins of *S. lincolnensis* are clindamycin-responsive thus the ABCF-mediated antibiotic signaling might be generally utilized tool of chemical communication.

**IMPORTANCE:** Resistance proteins are perceived as mechanisms protecting bacteria from the inhibitory effect of their produced antibiotic or antibiotics from competitors. Here, we report that antibiotic resistance proteins regulate lincomycin biosynthesis in response to subinhibitory concentrations of antibiotics. Particularly, we show the dual character of ABCF ATPase LmrC which confers antibiotic resistance and simultaneously transduces a signal from ribosome-bound antibiotic to gene expression, where the 5’ untranslated sequence upstream of its encoding gene functions as a primary antibiotic sensor. The ABCF-mediated antibiotic signaling can in principle function not only in the induction of antibiotic biosynthesis but in general in selective gene expression in response to any small molecules targeting the 50S ribosomal subunit, including clinically important antibiotics, to mediate intercellular antibiotic signaling and stress response induction. Moreover, the resistance-regulatory function of LmrC presented here for the first time unifies yet functionally inconsistent ABCF family involving the antibiotic resistance proteins and the translational regulators.

## INTRODUCTION

The genus *Streptomyces* and several other related genera of *Actinobacteria* (hereinafter streptomycetes) are filamentous soil bacteria characterized by a remarkably rich specialized metabolism. Specifically, the genomes of streptomycetes contain the highest proportion of biosynthetic gene clusters (BGCs) per Mb among all bacteria (1). The BGCs encode the biosynthesis of a wide arsenal of bioactive specialized metabolites, which have found their application in various areas, particularly in medicine. For instance, streptomycetes produce two-thirds of clinically used antibiotics of natural origin. However, the relevant biological roles of specialized metabolites in nature are still under debate. The current concept is that antibiotics are produced in response to cues from competitors to defend the habitats of their producers in natural competitive environments (2–5). These cues involve also ribosome-targeting antibiotics which at subinhibitory concentrations act as elicitors of secondary metabolism (6–8). However, antibiotic-sensing systems common for a group of functionally related but structurally distinct ribosome-targeting antibiotics have not been reported. Streptomycete-derived macrolide, ketolide, lincosamide, and streptogramin antibiotics target the peptidyl transferase center (PTC) of the 50S ribosomal subunit or its proximity (adjacent A- and P-sites or ribosomal exit tunnel). As a result, all these natural products interfere with proteosynthesis and inhibit bacterial cell-growth (for review see (9)). Therefore, apart from truly biosynthetic genes, BGCs of 50S ribosomal subunit-targeting antibiotics encode also resistance mechanisms for self-protection and regulation elements for timely and coordinated production. Typically, the BGC expression is directed by a pathway-specific regulator, which is activated by global and/or pleiotropic regulators (for review see (10). As for the resistance, one protein can be sufficient to protect the producing strain (11, 12). However, several mechanisms for self-resistance are often encoded in the BGCs for 50S ribosomal subunit-targeting antibiotics (13–17). It has been proposed that the expression of multiple resistance genes is regulated to optimize the self-protective resistance levels at different stages of growth or biosynthesis to minimize the fitness cost of the resistance expression (18) or to synchronize the resistance in sibling cells (19, 20). BGCs for 50S ribosomal subunit-targeting antibiotics often encode antibiotic resistance proteins of the ATP-binding cassette family F (ABCF), but mostly together with other resistance mechanisms. The ABCF proteins are cytosolic ATPases of the ABC superfamily, which confer resistance by ribosome protection (21) and not by an efflux as hypothesized for a long time. Apart from antibiotic resistance, bacterial ABCFs include also protein implicated in translational regulation (22, 23) but the function of the majority of bacterial ABCF remains unknown. Despite their distinct biological functions all characterized ABCFs to date act on the ribosome, and their common feature is an ATP-dependent modulation of the peptidyl transferase center (PTC) (reviewed in (24)). ABCFs are widely distributed in almost all bacteria, with the highest number per genome encoded in actinomycetes (8-11) (25). However, no actinobacterial antibiotic resistance ABCF was studied with the knowledge of its ribosome-associated function.

BGCs encoding structurally highly related natural lincosamide antibiotics celesticetin and lincomycin represent such a pair, mentioned above, with minimal vs. a complex set of resistance genes (12, 26). Based on our knowledge of biosynthesis of both lincosamide antibiotics and a comparative analysis of the respective BGCs (Figure 1a, for review, see (27)), there is only one non-biosynthetic gene, *ccr1*, in celesticetin BGC, coding for Ccr1 23S rRNA mono-methyltransferase, which is a self-protecting resistance protein modifying the ribosomal target of the producing strain. In contrast, there are four putative non-biosynthetic genes in lincomycin BGC: the resistance gene, *lmrB*, homologous to *ccr1*, a gene encoding the LmbU transcriptional regulator (28, 29), and two putative resistance genes coding for a transporter of the major facilitator family, LmrA, and an ABCF protein, LmrC.

**Figure 1.**
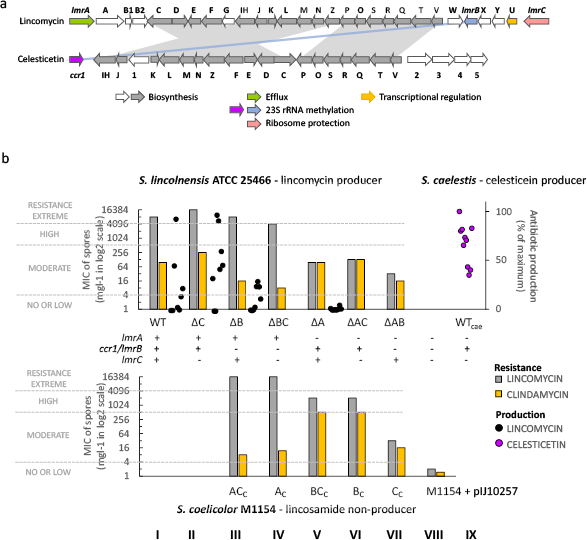
Contribution of LmrA, LmrB and/or LmrC resistance proteins to the self-protection and the production of lincomycin and celesticetin. **a**) Lincomycin and celesticetin biosynthetic gene clusters share eighteen biosynthetic genes encoding the common lincosamide scaffold (in grey) and *lmrB/ccr1* resistance genes (in blue and violet). Other structural genes encode the specific lincomycin and celesticetin biosynthetic steps (in white). The remaining three lincomycin BGC genes encode the transcriptional activator LmbUand resistance proteins LmrA and LmrC. **b**) Susceptibility of wild type (WT) and knock-out *S. lincolnensis* and *S. coelicolor* M1154 with an empty vector or with resistance genes under the control of the constitutive *ermEp* promoter (subscript c) – determined using spore suspensions; minimal inhibitory concentrations (MIC) were determined using spore suspensions. Mean MICs values (n≥3) are shown on a log2 scale, A – *lmrA,* B – *lmrB,* C-*lmrC.* Additional data (complementation with genes under the control of a putative natural promoter, resistance phenotypes determined using mycelia, the LmrC production levels from the natural and ermEp promoter) are available in Fig. S1 and S3b). Levels of lincosamides production of WT and single knock-out *S lincolnensis* and WT (n=8) and WT *S. caelestis* (n=10) are given as a percentage of the maximum production achieved.

In this study, we have shown that multiple resistance genes in the lincomycin BGC function together with the transcriptional regulator, LmbU, as a synchronized regulation-resistance unit. Although all Lmr proteins confer the resistance to lincosamides, only LmrA and partially LmrB affected overall lincomycin production. On the contrary, ribosome-operating LmrC ABCF ATPase is a key component of an antibiotic-induced cascade directing the onset of lincomycin biosynthesis through LmbU and in cooperation with LmrA and LmrB. This work represents the first reported antibiotic-driven activation of a BGC mediated by an ABCF resistance protein, *i.e*., a dual resistance-regulatory function of an ABCF protein.

## RESULTS

### LmrC antibiotic resistance protein is dispensable for resistance

To interpret the role of the three resistance proteins encoded in the lincomycin BGC, we first evaluated the contribution of the individual proteins to the resistance. Specifically, we knocked out *lmrA, lmrB*, and *lmrC* singly or in pairs in the lincomycin-producing *S. lincolnensis* wild type (WT) strain, and in addition, we complemented the genes under the control of a constitutive or natural promoter *in trans*(Fig. S1). Further, we constitutively expressed the genes in a lincosamide-sensitive *Streptomyces coelicolor* M1154 strain (30). Then, we evaluated the resistance phenotype of the WT, knock-out, and complemented strains by determining the minimal inhibitory concentration (MIC) of lincomycin and its derivative, clindamycin.

We revealed that all strains bearing *lmrA* including the strains with *lmrA* only (Fig. 1b columns I-IV) are highly or extremely resistant to lincomycin, while the resistance to lincomycin significantly decreases when *lmrA* is absent regardless of whether and which of the other two resistance genes are present. Interestingly, the resistance to clindamycin, generally a more efficient semisynthetic derivative of lincomycin, is different in this respect. Specifically, the majority of the tested strains are moderately resistant to clindamycin with no observed contribution of LmrA to the resistance (Fig. 1b – compare the columns differing in *lmrA* only: III vs. VII, I vs. V, II vs. VI). Therefore, we assume that LmrA, a transporter of the major facilitator family, is highly specific to lincomycin, but not clindamycin, and it ensures a sufficient self-resistance to the produced lincomycin on its own.

In contrast to the LmrA transporter, LmrB 23S rRNA monomethyltransferase confers the same or similar level of resistance (moderate or high) to both lincomycin and clindamycin as evidenced from the susceptibilities of the strains bearing *lmrB*, but not *lmrA* (Fig. 1b – columns V and VI).

The contribution of LmrB to the overall resistance is significant only when lincomycin-specific LmrA is not effective, i.e., for clindamycin (Fig. 1b – compare columns II vs. IV) or when LmrA is not present (Fig. 1b – compare columns V vs VII).

The last resistance protein, LmrC, confers moderate resistance to both lincomycin and clindamycin on its own (Fig. 1b – column VII). However, its contribution to the overall resistance is not considerable relative to neither LmrB (Fig. 1b – compare column V vs VI) nor LmrA (Fig. 1b – compare columns III vs. IV). The ΔC knock-out strain without *lmrC* shows even slightly increased lincosamide resistance compared to WT (Fig. 1b – compare columns I and II).

It is worth noting that the complementation of the knock-out strains under the control of the putative natural promoter restored the resistance phenotype of the WT except for *lmrB* (Fig. S1). In this case, the complementation had to be performed under the control of a constitutive promoter because *lmrB* is co-transcribed with three upstream genes as evidenced below. Interestingly, the constitutive *lmrB* expression resulted in higher resistance values compared to *lmrB* in its original genomic context (Supplementary Fig. 2). Further, the resistance of *S. lincolnensis* WT and knock-out strains were determined from spore suspension, which does not have to reflect the resistance of the mycelium during lincomycin production. Therefore, we determined MICs of *S. lincolnensis* deletion strains using mycelia from three different time points of the seed or production cultures (Fig. S1). On the whole, the data for mycelia comply with the data obtained for spores and show that resistance of the mycelium during production increased compared to the mycelium from the seed culture.

**Figure 2.**
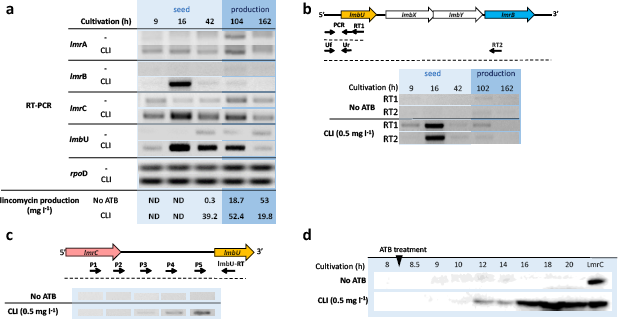
Clindamycin induces the expression of all resistance/regulation genes and hastens lincomycin production. **a,** RT-PCR detecting the presence of *lmbU, lmrA, lmrB* and *lmrC* transcripts on total RNA isolated from *S. lincolnensis* WT in indicated time points of cultivations without and with clindamycin supplementation (0.5 mg l^-1^, Fig. S7C), *rpoD* gene encodes the RNA polymerase sigma factor and was used as a control. **b,** RT PCR - *lmrB* is transcribed within the *lmbUXYlmrB* operon. **c,** RT PCR - *lmrC* and *lmbU* are transcribed independently **d,** Western blots showing LmrC protein levels in *S lincolnensis* WT without and with supplementation of clindamycin. Additional RT-PCR mapping of the lmrC-lmbU intergenic region is available in Fig. S2 and evaluation of the LmrC antibody is available in Fig. S3.

Apart from investigating the resistance phenotypes, we determined the amount of lincomycin produced by *S. lincolnensis* WT and single knock-out strains into the culture broth (Fig. 1b). The results support our conclusions drawn from the resistance of the strains. Specifically, only the strains with high or extreme resistance to lincomycin were able to produce lincomycin, i.e., the strains bearing both *lmrA* and *lmrB* (highest amount of produced lincomycin) and the strain bearing *lmrA* and not *lmrB* (up to 50% of the highest amount of produced lincomycin). Strain without *lmrA* failed to produce lincomycin, showing that the absence of LmrA prevents lincomycin biosynthesis likely because LmrB, under the control of the natural promoter, is not available at the required time and LmrC does not ensure sufficient protection of the cell. The dispensability of LmrC for the overall resistance documented above complies with the comparable lincomycin production of ΔC vs. WT strains. In addition, the production of lincomycin significantly fluctuated (Fig. 1b). This observation could be explained by a more complex regulation-resistance system (LmbU, LmrA, LmrB, LmrC) encoded within the lincomycin BGC compared to the BGC of another lincosamide, celesticetin (only Ccr1 from the same family as LmrB) (Fig. 1a). Indeed, the fluctuation of celesticetin produced by *S. caelestis* in parallel cultures is not as pronounced as that of lincomycin produced by *S. lincolnensis.* Moreover, *S caelestis* never failed to produce celesticetin, while *S lincolnensis* failed to produce lincomycin in several parallel WT cultures (Fig. 1b).

### Expression of *lmrA, lmrB, lmrC,* and *lmbU* is induced by clindamycin

Given our hypothesis of the complex regulation-resistance system of lincomycin production, we were wondering whether the expression of any of *lmbU, lmrA, lmrB,* and *lmrC* genes could be affected by the produced antibiotic. Therefore, we cultured *S lincolnensis* WT and divided the culture before the onset of lincomycin biosynthesis into two parallels, one of which was supplemented with clindamycin at a subinhibitory concentration (Fig. 2a). At several time points, we semi-quantitatively monitored the expression of the respective genes by reverse transcription PCR (RT-PCR). The supplementation with clindamycin allowed us to distinguish between the lincosamide used to study its effect on the gene expression and the lincosamide produced by the strain, which we determined by ultra-high-performance liquid chromatography.

The results summarized in Fig. 2a show a dramatic effect of the supplementation with clindamycin on the expression of all the studied genes, *lmbU, lmrB, lmrA,* and *lmrC*, while no effect was observed on the *rpoD* control. The most pronounced was the clindamycin-induced expression of *lmbU* and *lmrB.* These transcripts were detected at earlier time points compared to the untreated cultures. In agreement with this observation, also the onset of lincomycin production is shifted towards an earlier time of cultivation in the culture supplemented with clindamycin (Fig. 2a). Similar profiles of *lmbU* and *lmrB* transcripts indicate that *lmrB* is organized in the operon with the regulation and biosynthetic genes, *lmbUXY*. Indeed, amplification of *lmbU* from the 1st DNA strand synthesized using a primer specific to *lmrB*, confirms that the expression of *lmrB* is directly coupled with that of *lmbU* and the two biosynthetic genes, *lmbX* and *lmbY* (Fig. 2b). In an analogous mapping strategy, we confirmed that the *lmbUXYlmrB* operon is transcribed independently of the upstream *lmrC* gene (Fig. 2c, Fig. S2) and thus also their induction by clindamycin on the level of transcription is independent. In contrast to *lmbU, lmrB,* and *lmrA* transcripts, *lmrC* was detected in all samples, but at higher levels in the cultures supplemented with clindamycin. We subsequently confirmed that *lmrC* expression is clindamycin-responsive on the protein level by monitoring LmrC using an LmrC-specific antibody in the cells from *S. lincolnensis* WT cultures without and with clindamycin supplementation (Fig. 2d).

### LmrC is essential for the antibiotic-induced onset of lincomycin production

Given the newly defined function of ABCF proteins as modulators of ribosomal PTC, the onset of lincomycin production in response to antibiotics might be regulated by LmrC. To uncover the role of LmrC, we performed comparative mass spectrometry proteomic analysis of the mycelia of *S lincolnensis* WT, WT+Cc, and ΔC strains grown in the absence or presence of clindamycin. As shown in Fig. 3a, the induction of expression of lincomycin BGC proteins by clindamycin is substantially higher in WT+Cc compared to WT, while the clindamycin induction of the gene expression is not observed in the ΔC strain. These experiments suggest that LmrC is required for the induction of gene expression triggered by clindamycin. To confirm that LmrC is essential for transduction of the antibiotic signal to the expression of lincomycin BGC, we quantified the transcripts of *lmrC, lmbU*, and a biosynthetic gene *lmbN*, which is not under the direct control of *lmbU* in *S lincolnensis* WT and ΔC cultured without and with clindamycin in a time point before the lincomycin BGC expression. As shown in Fig. 3b, clindamycin induced *lmrC, lmbU*, and *lmbN* transcription in WT, but not in the *lmrC*-deficient ΔC knock-out strain. It is noteworthy that the observed low-level constitutive transcription of *lmbU* in the ΔC strain can be explained by the insertion of the apramycin cassette (Fig. S2b), causing a polar effect. This phenomenon explains the increased production of proteins in ΔC (Fig. 3a). However, it is important for our reasoning that neither the protein production nor the *lmbU* transcription in the ΔC strain is affected by clindamycin. The ABCF family proteins generally exhibit an ATPase activity, which is required for the protein function (25). We were, therefore, wondering whether LmrC has to be a functional protein capable of the ATPase activity to induce the gene expression. Hence, we complemented the ΔC knock-out strain with *lmrC* or *lmrC_EQ12_* expressed from a theophylline-inducible plasmid (C_i_). Overproduction of functional LmrC resulted in the expression of *lmbU* and *lmbN*, while the overproduction of the ATPase-deficient LmrC_EQ12_ mutant did not have this effect (Fig. 3c). Notably, the expression of *lmbU* and *lmbN* mediated by the overproduction of LmrC was achieved without the supplementation with clindamycin, and a similar phenomenon was observed on the protein level when LmrC was produced constitutively in WT (compare WT and WT+C_C_ without clindamycin supplementation in Fig. 3a). The induction by clindamycin is observable also on the lincomycin production level in WT, and ΔB but not in ΔC knock-out strain where higher production levels were independent of clindamycin supplementation (Fig. 3d). These results demonstrate that clindamycin induces the production of LmrC, which in turn induces the production of LmbU, which is a known activator of lincomycin biosynthesis (28).

**Figure 3.**
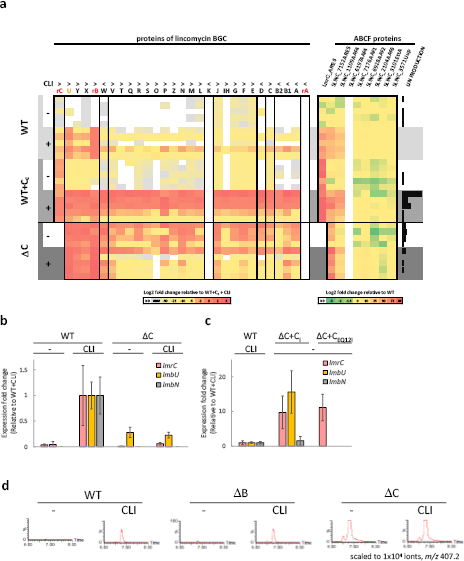
ATPase active LmrC transduces the antibiotic signal to trigger lincomycin production in response to exogenously added clindamycin by inducing the transcription of *lmbU.* **a,** Relative abundance of lincomycin biosynthetic gene cluster (BGC)-encoded proteins and ABCF proteins in *S. lincolnensis* WT, WT constitutively expressing *lmrC*(WT+Cc) and the *lmrC* knock out strain (ΔC) cultivated for 40 h with or without clindamycin (CLI) (Supplementary Fig. 3B). The expression ratios are defined for the lincomycin BGC-encoded proteins as the fold change relative to the median expression levels in the clindamycin-treated samples of the WT+Cc strain and for ABCF proteins as the fold change relative to the median expression levels in the untreated WT. A comparison of growth with and without clindamycin supplementation is available in Fig. S4b **b,** qRT-PCR analysis (n=4) of *lmrC, lmbU,* and *lmbN* transcripts in the WT and ΔC in a 16h seed culture (Fig. S7C). The data are expressed relative to WT cultured with clindamycin, error bars indicate standard deviation. **c,** qRT-PCR analysis of *lmrC, lmbU,* and *lmbN*transcripts in ΔC with theophylline-inducible production of the unmutated LmrC (ΔC+Ci) and its ATPase-deficient mutant LmrC_EQ12_ (ΔC+C_EQ12i_). Cultivation and data processing are identical to those in b. Proof that LmrC_EQ12_ does not affect the growth is available in Fig. S4a. **d,** Lincomycin produced into the media of WT (n=6), lmrB knock-out strain (ΔB, n=2), and ΔC (n=4) from 42 h seed culture with and without clindamycin supplementation (Fig. S7b). Representative extracted ion chromatograms of lincomycin, which elutes in 6.9 min are shown. All chromatograms are available in Fig. S5.

### LS_A_P group antibiotics induce LmrC production *via* ribosome-mediated transcriptional attenuation

To clarify the mechanism of induction of *lmrC* expression, we first tested whether the inducing antibiotics are limited only to clindamycin. For this purpose, we treated *S. lincolnensis* WT cells with a range of ribosome-targeting antibiotics and a cell wall-targeting carbenicillin, and we detected LmrC protein levels using a specific antibody. In addition to clindamycin, only lincomycin (lincosamide group), pristinamycin IIA (streptogramin A group), and with lower efficiency tiamulin (pleuromutilin group) induced LmrC production (Fig. 4a). Interestingly, all these compounds belong to the PTC-targeting antibiotics of the lincosamide-streptogramin A-pleuromutilin (LS_A_P) group (Fig. 4a), to which antibiotic resistance ABCF proteins, exemplified by Vga(A)_LC_, confer resistance in clinical isolates (21, 31). It suggests that the regulation of LmrC production is coupled to its resistance function that is enabled by dislocating the antibiotic from its specific overlapping binding sites within the PTC.

**Figure 4.**
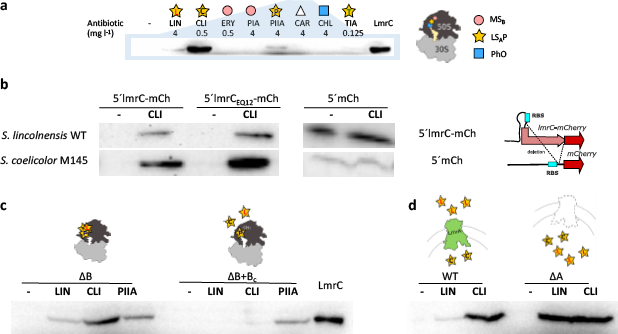
LmrC expression is induced by LS_A_P-group of antibiotics *via* ribosome-mediated attenuation and dampened by LmrA and LmrB Western blot analysis of LmrC production in 16h seed culture mycelium uninduced or induced by antibiotics (Fig. S7c). Protein production was detected either using an LmrC antibody (a, c, d), or mCherry-specific antibody (b). **a,** LmrC levels in the WT 8 h after addition of antibiotic: LIN – lincomycin, CLI – clindamycin, PIIA – pristinamycin IIA, TIA – tiamulin, ERY – erythromycin, PIA – pristinamycin IA, CHL – chloramphenicol, and CAR - carbenicillin. Schematic illustration of the overlapping binding sites of the antibiotic groups (represented by colored symbols) defined by resistance phenotypes conferred by ARE: LS_A_P – lincosamides, streptogramins A, pleuromutilins; _MSB_ – macrolides, streptogramin B, PhO – phenicol, oxazolidinones. **b,** mCherry levels in *S. lincolnensis* WT and *S. coelicolor* M145 carrying plasmids 5’lmrC-mCh or 5’lmrC_EQ12_-mCh, in which region encoding a 405 bp *lmrC* upstream sequence and the full-length *lmrC* gene (or its ATPase deficient mutant *lmrC*_EQ12_), were fused to mCherry, and the plasmid *5’mCh,* in which an identical 405 bp upstream region, was inserted in front of mCherry without the *lmrC* gene; in the latter case, the putative terminator hairpin, which partially overlaps the *lmrC* gene, cannot be formed. **c,** LmrC levels in ΔB compared to those of ΔB+Bc with constitutive expression of lmrB are lower in the case of lincomycin and clindamycin induction because these cannot bind to the ribosomes methylated by LmrB; pristinamycin IIA can bind to the methylated ribosome and its induction of lmrC expression is therefore not affected. Additional data showing LmrB-mediated feedback on the lincomycin biosynthesis level are available in Fig. S5; **d,** LmrC levels in the WT compared to ΔA with the lincomycin-specific exporter LmrA deleted; LmrA reduces the intracellular concentration of lincomycin affecting thus the induction of *lmrC* expression.

Previous work has shown that the expression of antibiotic resistance ABCF proteins is controlled in response to ribosome-targeting antibiotics through ribosome-mediated attenuation (32–34) and could be shaped by the resistance activity of the protein (33). To prove whether the attenuation mechanism is involved in the control of *lmrC* expression, we performed an *in silico* analysis of the *lmrC* 5’ UTR which revealed features characteristic of this regulatory mechanism (35) and mapped the length of the transcript by RT-PCR. Specifically, we identified 5’ UTR of about 230 nucleotides with a putative promoter, premature terminator with the ability to form alternative anti-terminator conformations, and a uORF, with a strong ribosome binding site (RBS) (Fig. S6).

To determine whether the putative premature terminator contributes to the regulation of *lmrC* expression, we prepared a fusion construct of a mCherry reporter with the *lmrC* upstream region (including its promoter) inserted either with the full-length *lmrC* (5’ lmrC-mCh), inactive mutant lmrC_EQ12_ (5’ lmrC_EQ12_-mCh) or alone (5’ mCh). We introduced the constructs into *S. lincolnensis* WT and *S. coelicolor* M145 strains, cultured the strains without and with clindamycin, and determined mCherry levels (Fig. 4b). In strains with plasmids containing an intact premature terminator, the mCherry-specific band was detected only in the presence of clindamycin and the activity of LmrC had no or minor effect on the induction (compare 5’ lmrC-mCh and 5’ lmrC_EQ12_-mCh in Fig, 4b). In contrast, strains with the reporter plasmid (i.e., the construct without the *lmrC* sequence), in which the terminator hairpin cannot be formed, produced mCherry regardless of clindamycin (Fig. 4b). The antibiotic-mediated control of *lmrC* expression thus occurs *via* a formation of the premature terminator structure, which prevents *lmrC* expression in the absence of an antibiotic. LS_A_P antibiotics, presumably if bound to the PTC, trigger the shift from the anti-terminator to the terminator conformation, enabling *lmrC*transcription (Fig. S5c). In addition, the induction is not dependent on another *S lincolnensis* gene because the induction was functional even in the *S coeilcolor* heterologous host (Fig. 4b). The *lmrC* transcript, specifically its attenuator, is thus the primary sensor of the antibiotic-LmrC-LmbU signaling cascade for lincomycin biosynthesis.

### LmrA and LmrB provide a negative feedback loop to the cascade by reducing *lmrC* expression

We have shown that the expression of *lmrC* responds to the presence of LS_A_P antibiotics (Fig. 4a). The other lincomycin BGC-encoded resistance proteins, LmrA and LmrB, influence the activity of some of these antibiotics; therefore, we investigated whether they can affect the whole cascade by dampening *lmrC* expression. First, we evaluated LmrC protein levels in *S. lincolnesis* ΔB+B with the constitutive overproduction of the LmrB methyltransferase. As shown in Fig. 5a, LmrC production no longer responded to lincomycin and clindamycin that do not bind to ribosomes methylated by LmrB, but it remained responsive to the treatment with pristinamycin IIA (streptogramin A group), which can bind to methylated ribosomes (36). This experiment also demonstrates that the binding of LS_A_P antibiotics to the ribosome is the prerequisite for the induction of *lmrC* expression thereby further approving the ribosome-mediated attenuation mechanism.

**Figure 5.**
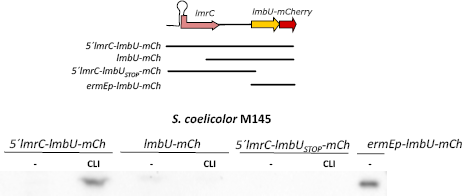
The antibiotic signaling cascade hastening the onset of lincomycin production consists in its basic form only of *lmrC* and *lmbU* genes Western blot analysis of LmbU-mCherry levels in 16h seed culture mycelium (Fig. S7c) uninduced or induced by clindamycin (CLI, 0.03 mg l^-1^). LmbU-mCherry production was detected by mCherry-specific antibody in *S. coelicolor* M145 carrying plasmid 5’lmrC-lmbU-mCh, lmbU-mCh, and plasmid 5’lmrC-lmbU_ST0P__mCh as the negative control.

Since the expression of *lmrB* is in the *lmbUXYlmrB* operon with the *lmbX* and *lmbY* biosynthetic proteins (Fig. 2b), LmrB is an ideal candidate to provide a feedback loop of the cascade. Indeed, pronounced induction of lincomycin production was apparent in ΔB compared to WT while in the ΔB+B strain, the constitutive *lmrB* expression dampened the onset of lincomycin biosynthesis (Fig. S5). Next, we evaluated LmrC protein levels in *S lincolnensis* WT after the addition of lincomycin or clindamycin. The considerably higher level of LmrC in the clindamycin-induced compared to lincomycin-induced WT cells (Fig. 5b) complies with LmrA specifically transporting lincomycin, but not clindamycin out of the cell. The low intracellular level of lincomycin induces *lmrC* expression less effectively than the high level of clindamycin. This was supported by an analogous experiment with LmrA-deficient *S. lincolnensis* ΔA strain, in which LmrC expression was induced by lincomycin and clindamycin at a comparable level (Fig. 5b). It shows that LmrA specifically dampens LmrC expression induced by lincomycin by reducing its intracellular concentration. Since both LmrB and LmrA reduce *lmrC*expression in response to antibiotics, we propose that alongside their resistance function they also serve as a negative feedback loop to the antibiotic-LmrC-LmbU signaling cascade of lincomycin biosynthesis.

### Antibiotic-LmrC-LmbU signaling cascade is independent of other *S. lincolnensis* regulatory elements

Several recent studies described regulators of lincomycin biosynthesis encoded outside the BGC in the *S. lincolnensis* genome (37–40). Some of these regulators might be involved in the antibiotic-induced onset of lincomycin production. To rule out this hypothesis, we cloned the lincomycin BGC region starting upstream of *lmrC* and ending with *lmbU* translationally fused with mCherry reporter (5’lmrC-lmbU-mCh) and introduced into *S. coelicolor* M145. A truncated version of the construct without the 5’half of *lmrC*(lmbU-mCh) and a construct with a non-sense mutation in *lmbU-mCh* (5’lmrC-lmbU_ST0P__mCh) were used as negative controls and lmbU-mCherry expressed from constitutive ermEp promoter (lmbU-mChc) as a positive control (Fig. 5a). Slight or no production of LmbU-mCherry was detected in both uninduced 5’lmrC-lmbU-mCh and lmbU-mCh, but only the 5’lmrC-lmbU-mCh construct, producing LmrC, increases the expression of LmbU-mCherry after the addition of clindamycin. These data show that the cascade triggering the onset of lincomycin production consists in its basic form only of *lmrC* and *lmbU* genes and that none additional elements from *S. lincolnensis* are required.

### LmrC is coproduced with seven other ABCFs, two of which are responsive to a lincosamide

In addition to resistance, the LmrC ABCF protein has a regulatory function, which transduces an antibiotic signal to activate lincomycin biosynthesis in *S. lincolnensis.* Comprehensive phylogenetic analysis classified 30 subfamilies of bacterial ABCF proteins (25). Four subfamilies (Uup, Etta, YdiF, and YbiT) have a broad distribution while others including subfamilies with the resistance function (ARE1-7) are taxon-specific. *Actinobacteria* is a phylum with the highest number of ABCFs including seven subfamilies specific to this taxon (AAF1-6, ARE4-5). In addition to LmrC, which belongs to the ARE5 subfamily, the genome of *S. lincolnensis* encodes eight ABCF proteins - three of them with a putative resistance activity (ARE5 encoded by SLINC_7152 and two AAF4 encoded by SLINC_1109 and SLINC_6197). We were wondering whether some of these resistance proteins are induced by clindamycin and thus they could have an antibiotic-responsive regulatory function. We used the same mass spectrometry proteomics data set as for the comparative analysis of lincomycin biosynthetic proteins to visualize the abundance of ABCF proteins in *S. lincolnensis* WT, WT+Cc, and Δ_c_ strains grown in the absence or presence of clindamycin. As shown in Fig. 3a, all but one ABCF proteins were present in all samples but only two out of three putative antibiotic-resistance ABCF proteins were substantially induced by clindamycin independently on the LmrC presence. The third putative resistance ABCF protein was not detected in any of the samples. Considering that the putative resistance function of these clindamycin-responsive ABCF proteins is abundant, they could have a regulatory function similar to LmrC.

## DISCUSSION

Antibiotic resistance proteins associated with BGCs have traditionally been perceived as a means of self-protecting mechanisms. Their role in the regulation of production of antibiotic metabolites has been proposed, (41) but not demonstrated. In this study, we characterized an LS_A_P-antibiotic-driven signaling cascade for the activation of the onset of lincomycin biosynthesis, in which an antibiotic resistance protein, LmrC, from the ARE5 subfamily of ABCF proteins is the key signal-transducing element (Fig. 6a). The mechanism lies in the induction of *lmrC* transcription by a ribosome-mediated attenuation, which means that *lmrC,* specifically its attenuator-forming upstream 5’UTR transcript, is a sensor of LS_A_P antibiotics. The ribosome-mediated attenuation is a common mechanism of regulation of antibiotic resistance ABCF genes in Firmicutes (32–34, 42). However, in this work, it is for the first time described to function as a sensor of the signaling cascade. The major novelty of this cascade lies in the dual antibiotic resistance and regulatory function of the ABCF protein, LmrC, which transduces the antibiotic signal to the expression of LmbU, which promotes lincomycin biosynthesis. In addition, we have shown that another two lincomycin BGC-encoded resistance proteins, LmrB and LmrA, affect the cascade by dampening the LS_A_P antibiotic-induced expression of *lmrC.* We assume that LmrB, by its position in the *lmbUXYlmrB* operon with two biosynthetic genes, mediates a direct negative-feedback loop of the cascade. LmrA transporter links lincomycin biosynthesis to primary metabolic pathways as it is regulated by the GlnR global regulator (43). LmrA seems to be the most important for lincomycin production, without it, lincomycin biosynthesis is remarkably suppressed. Further, LmrA, as a lincomycin-specific transporter, desensitizes the cascade only to lincomycin, which may prevent reactivation of the biosynthesis by its product when it is no longer desirable. In addition, the active export of lincomycin contributes to the propagation of the antibiotic within the population. The last component of the regulation cascade, LmbU, is a transcriptional regulator of the newly proposed LmbU-family (28). The *lmbU* gene was evolutionarily accepted together with genes encoding the unusual 4-alkyl-L-proline precursor, (27) a common building block of lincomycin and other natural products of streptomycetes (44–46) but not the proteinogenic L-proline incorporating celesticetin, in which BGC *lmbU* homolog is logically missing.

**Figure 6.**
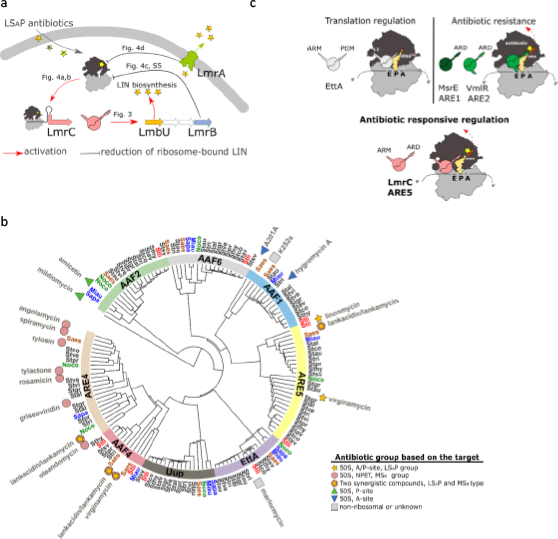
ABCF proteins encoded in biosynthetic gene clusters are putative regulators of antibiotic production in response to antibiotics that share ribosomal binding site. **a,** Scheme of the antibiotic-LmrC-LmbU signaling cascade identified in this study. The production of the LmrC protein is induced by ribosomal-bound LS_A_P antibiotics *via* a ribosome-mediated attenuation mechanism, and it is coordinated with the LmrB and LmrA resistance proteins, which individually reduce the amount of ribosome-bound antibiotic. The LmrC then transduces the antibiotic signal from the ribosome to the transcription of *lmbU.* The LmbU transcriptional regulator activates the expression of the subordinate biosynthetic genes (28). **b,** Phylogenetic tree of ABCF proteins from 14 representative streptomycetes genomes and ABCF proteins from previously characterized BGCs. ABCF proteins from characterized BGCs are marked with the name of produced antibiotic: symbols in the legend indicate the antibiotic group. The genomic ABCF protein sequences were taken from previously published data (25). A list of streptomycetes genomes and BGCs is available in Table S1. **c,** LmrC domain architecture combines features of resistance and regulatory ABCF proteins. The presence of the arm domain, resembling the ABCF translation regulator EttA, indicates the regulatory function of LmrC, while the antibiotic resistance domain (ARD) is shared with other structurally characterized ABCF resistance proteins. Peptidyl tRNA interaction motif (PtIM) in EttA and ARD structural motifs refer to a linker that separates two ATP binding domains. ARD domain is significantly longer than PtIM allowing direct interaction with PTC.

The regulatory pair of LmbU and LmrC is unique to lincomycin BGC – no other known BGC encodes a LmbU-family regulator together with an ABCF protein. On the other hand, BGC-associated ABCF proteins are almost exclusively present in the BGCs for PTC-targeting antibiotics (Fig. 6b, Table S1). Most of these BGCs encode additional resistance determinants and pathway-specific transcriptional regulators; however, none is homologous to LmbU (Table S1). We assume that the BGC-encoded ABCF proteins employ transcriptional regulators of diverse families to form a signaling cascade for the activation of biosynthesis of the ribosome-binding antimicrobials. We can hypothesize that the absence of a transcriptional regulator in the celesticetin BGC did not allow establish a resistance regulation unit analogous to LmrC-LmbU pair in the lincomycin biosynthesis.

It was previously shown that LmbU directly activates only 4-alkyl-L-proline biosynthesis-encoding part of the lincomycin BGC (28), which is also evident from our proteomic data (Fig. 3a). Recently, a new regulator of lincomycin BGC, AdpAlin, which activates the entire lincomycin BGC independently on the external lincosamide and thus appears to be the master regulator, has been described (37). What would then be the purpose of the ABCF protein, LmrC, and its involvement in the here discovered LmrC-LmbU signaling cascade? It may be solely the alternative activation of lincomycin BGC in response to the presence of lincomycin towards synchronizing the specialized metabolism within a population of siblings. Such an advanced level of regulation of antibiotic production mediated by BGC-associated ABCFs could provide a strong competitive advantage in the natural environment. This hypothesis is supported by the fact that all except one BGC-encoded ABCF proteins are encoded in known BGCs for PTC-targeting antibiotics (Fig. 6b). Moreover, the type of ABCF protein strictly follows the exact binding site of the antibiotic produced, even to the extent that BGCs for synergistic pairs of structurally different compounds such as streptogramins (47, 48), encode ABCF proteins of two different subfamilies, ARE5 together with either ARE4 or AAF4, reflecting two distinct binding sites of these compounds (Fig. 6b). Besides, analogous ABCF-mediated antibiotic-signaling cascades can contribute to the cooperative competition, allowing strains with BGCs for functionally similar 50S ribosomal subunit-targeting antibiotics to synchronize the production and thus to improve the competitiveness over others (49). In the case of the LmrC-LmbU cascade, the cooperating strains would represent producers of antibiotics inducing *lmrC* expression, *i.e*., LS_A_P antibiotics and other molecules binding to the same site on the ribosome. In support of this concept, a recent study showed that antibiotic production is more likely to be induced by closely related strains or strains sharing BGCs (5). These observations also imply that the antibiotic-induced onset of antibiotic production is a relatively widespread but mostly undetected phenomenon because the production in the presence of cognate or similar antibiotics is not usually examined.

The induction of specialized metabolism by antibiotics targeting the 50S subunit of the ribosome has been described previously (6, 7, 50), but this is the first time when the mechanism of antibiotic sensing and signal transduction has been described. The detection of antibiotic by a ribosome *via* 5’UTR attenuator upstream of ABCF encoding gene differ fundamentally from known antibiotic signaling cascades of other types of antibiotics, in which the antibiotic or a biosynthetic intermediate are detected regardless of their mode of action - typically by direct binding to a transcription factor or its cognate receptor (51). The role of antibiotic resistance ABCF proteins in the transduction of the antibiotic signal to the transcriptional regulator is also unprecedented, but perhaps not so surprising. LmrC, as well as other ABCF proteins implicated in antibiotic resistance in streptomycetes, show structural similarities with both the antibiotic resistance ABCF proteins MsrE and VmlR as well as with the translational regulator EttA (Fig. 6c): LmrC shares with antibiotic resistance protein the antibiotic resistance domain, ARD, which interacts with the PTC to dislodge the antibiotic from the ribosome (31, 52, 53). ARD is present in the majority of ARE ABCF proteins but not in other types of ABCFs, and thus it reflects their antibiotic resistance function (25). Moreover, LmrC has also the arm domain, which is absent in VmlR and MsrE but is present in EttA (22). The arm domain restricts in EttA the ribosome dynamics in response to a lack of available ATP (Boël et al., 2014). Since ATP hydrolysis was required for the antibiotic signal transduction to LmbU (Fig. 3c), LmrC might integrate the antibiotic signal and, analogously to EttA, the information about the energetic status of the cell. Further research will be, however, needed to determine whether all structurally similar ABCF proteins have regulatory rather than resistance function regardless of whether they are in the BGC of the respective antibiotic. The fact that, in addition to LmrC, another two ABCF proteins were induced by clindamycin, regardless of the presence of LmrC, is a strong indication of their antibiotic-responsive regulatory function.

The signaling pathway described herein, in which the antibiotic signal is sensed and transduced by a dual, resistance and regulatory ABCF protein, and tuned by two other resistance proteins, informs the need to reconsider the role of resistance proteins in biosynthetic pathways as purely self-protective mechanisms. In particular, the discovery of the dual function of LmrC resistance protein launches the breakpoint in overall perception of bacterial ABCF proteins as yet functionally inconsistent group involving the proteins conferring resistance to 50S ribosomal subunit-targeting antibiotics and the ribosomal regulators (25, 54). The LmrC is the first bacterial ABCF with demonstrated dual, resistance and regulatory function, unifying thus this yet split group. Besides, given the number of small molecules targeting the 50S ribosomal subunit and the number of bacterial ABCF encoded by soil bacteria from the Terrabacteria group, which includes Firmicutes and Actinobacteria with the highest number of ABCF per genome, ABCF-mediated signaling could be one of the most important tools of chemical communication in general.

## MATERIALS AND METHODS

Bacterial strains and growth conditions. The strains, plasmids, and oligonucleotides used in this study are listed in Table S2. *Streptomyces* strains were grown at 30°C on solid MS medium (55) (Mannitol Soya flour medium), DNA (2.3% Difco nutrient agar) and MH agar (1.5% agar in Mueller Hinton broth, purchased from Oxoid) or in liquid YPM2 (0.4% Yeast extract, 0.5% Peptone, 1% Malt extract, pH 7.2) or AVM (56) media. Spore suspensions of *Streptomyces* spp. were prepared and germinated for 3h in 2 × GM (1% yeast extract, 1% casaminoacids, 0.01 M CaCl_2_) according to protocols published in Practical Streptomyces Genetics (57). For the selection of exconjugants antibiotics were added into the cultivation media at the following concentrations: apramycin, 50 mg l^-1^; kanamycin, 50 mg l^-1^; carbenicillin, 100 mg l^-1^; chloramphenicol, 25 mg l^-1^; nalidixic acid, 25 mg l^-1^; and hygromycin at 100 mg l^-1^ in agar plates or at 80 mg l^-1^ (*E colt*) or 40 mg l^-1^ (*Streptomyces*) in liquid media. For lincomycin and celesticetin production and proteomic analysis, *S. lincolnensis* and its derived mutants or *S. caelestis* spores were germinated and inoculated into 50 ml of YPM2 (to reach OD450 0.03) in 250 ml baffled flasks and cultivated on an orbital shaker for 42 h at 30°C and 200 rpm. 2.5 ml of the whole YPM2 preculture was used to inoculate 47.5 ml of fresh AVM broth (56) followed by cultivation in 250 ml baffled flasks for 120 h at 30°C and 200 rpmFor RT-PCR, qPCR analyses and western blot analyses, the spores were germinated, inoculated 50 ml falcon tubes containing 20 ml YPM2 media (to reach OD450 0,03) and grown in on an orbital shaker (8 h, 30°C, 200 rpm) prior to induction. Then, the cultures were induced with an antibiotic indicated, and the cultivation continued for an additional 8 h (Fig. S7c).

### Construction of knock-out strains

The *S. lincolnensis* ΔA (BN3024), *S. lincolnensis* ΔB (BN3002) and *S. lincolnensis* ΔC (BN3001) mutants were constructed by replacing the entire coding sequence of the target gene with a cassette (773 or 775 (58)) carrying the apramycin resistance gene (*aac(3)IV*) and oriT of RK2, using the PCR-targeting method (59). Oligonucleotides used for gene deletion and verification of the deletion are listed in Table S2. PCR targeting was applied to the cosmid LK6 (26), which contains the entire lincomycin biosynthetic cluster. After conjugation of mutated cosmids into *S. lincolnensis,* kanamycin-sensitive (Kan ^S^), apramycin-resistant (Apra ^R^) double-crossover mutants with target genes replaced by the *aac(3)IV-oriT* cassette were confirmed by PCR amplification. For the construction of the *S. lincolnensis* ΔAB (BN3021), *S. lincolnensis* ΔAC (BN3018), and *S. lincolnensis* ΔBC (BN3008) double mutants, the inactivation cassettes 773 in *S. lincolnensis* ΔB (BN3002) and *S. lincolnensis* ΔC (BN3001) single mutants were replaced by an unmarked in-frame deletion, obtained by FLP-mediated excision of the disruption cassette (58). The second gene to be deleted was replaced with the cassette 775 according to the same protocol as for single knock-out strains. Knock-out strains were verified by PCR and Southern blot analysis. The scheme of the orientation of inactivation cassettes in all knock-out strains is available in Fig. S1.

### Construction of vectors for constitutive and inducible expression

Vectors pLV012, pLV017, and pLV013 were used to express resistance genes, each under its promoter. pLV012 was constructed by insertion of a 2,946 kb EcoRV fragment of LK6 containing *lmrA* with its 1,330 bp upstream sequence ligated into the EcoRV site of pMS81. pLV017 contains a 3,362 kb DNA fragment covering the gene *lmrC and its* 1,281 bp upstream region obtained by EcoRV and BglII digestion from cosmid LK6. pLV013 contains a 958 bp ApaI fragment of LK6 containing the *lmrB* gene with its 86 bp upstream region. If needed, DNA fragments were blunted by DNA polymerase I or the large Klenow fragment and then ligated into the EcoRV site of pMS81. For constitutive expression, *lmrA, lmrB,* and *lmrC* were PCR-amplified from LK6 and ligated under the *ermEp* promoter of pIJ10257 (60), *i.e*. between the NdeI and HindIII restriction sites (constructs pLV026, pLV030, and pLV018, genes marked with subscript c). For coexpression of *lmrA* or *lmrB* with *lmrC* in the heterologous host, *lmrC*with the *ermEp* promoter was cloned between the BglII and EcoRI restriction sites of the *PtipA* expression vector pIJ6902 (61) (construct pLV028, *lmrC* marked with subscript c2). Plasmid pLV025, for expression of the lmrC_EQ12_ mutant, was prepared by site-directed mutagenesis, where two mutations were introduced into the *lmrC* coding sequence: glutamate 167 was replaced with a codon for glutamine, and the parallel codon for glutamate 495 was replaced with a codon for glutamine using oligonucleotides Lmr(C)G499C-For, Lmr(C)G499C-Rev, Lmr(C)G1453-For and Lmr(C)G1453-Rev and pLV20 plasmid as template. Plasmids were amplified via whole-plasmid PCR, and DpnI was used to digest the methylated parental DNA. The resulting constructs were sequenced to verify that the expected mutations had been made. Inducible expression was performed using the pIJ10257 vector, which encodes a theophylline-dependent riboswitch (62). The riboswitch region was introduced into vectors pLV018 and pLV025 using whole-plasmid PCR with oligonucleotides RBSW_E*_lmrC F and pIJ_lmrC _R and subsequent DpnI treatment, T4 polynucleotide kinase 5’phosphorylation and plasmid recircularization, yielding vectors pGBN025 (lmrCi), and pGBN030 (lmrC_EQ12is_). The constructs pGBN054 (for constitutive fusion expression of *lmrC-mCherry* (*lmrC-mCh_c_*) was prepared using the SLICE cloning method (63). In pGBN054, we prepared two inserts – specifically, *lmrC* (PCR amplification from template pLV018, oligonucleotides pIJ_LmrC_lin_F and pIJ_LmrC_PGGGS_lin_R) and *mCherry* (PCR amplification from template mCherry (64), primers mCherry_pIJ_LmrC_F and mCherry_pIJ_LmrC_R), treated them with DpnI and cloned them into pIJ10257 under the control of the *ermEp* promoter. The pMK037 construct, bearing the 405 bp long *lmrC* upstream region cloned in front of *mCherry* (5’C-mCh), was also prepared using SLICE cloning. The PCR amplicon from the pLV017 template (oligonucleotides LmrCp-F, lmrCp-mCherry-R, DpnI treatment) was cloned into pGBN054 that had been linearized with NdeI and Bsu36I. Then, 5’C-*lmrC-mCh* (pMK038) was prepared by SLICE cloning of the 405 bp *lmrC* upstream region (PCR amplification from template pLV017, oligonucleotides LmrCp-F, lmrCp-lmrCr) into vector pGBN054 linearized with NdeI and Bsu36I. Plasmids 5’-*lmrC-lmbU-mCh* (pGBN062), *lmbU-mCh* (pGBN064) and 5’-*lmrC-lmbUsT0P-mCh* (pGBN065) were prepared by SLICE cloning method: vector pGBN054 was linearised via whole plasmid PCR (oligonucleotides pGBN054_F2 and pGBN054_R) and ligated with inserts 5’*-lmrC-lmbU* (primers pGBN054_lmrCU_F1 and pGBN054_wo_int_RBS_lmrCU_R, template pMK001) in pGBN062 and pGBN065; *lmbU* (primers pGBN054_lmrCU_F2 and pGBN054_wo_int_RBS_lmrCU_R, template pMK001) for pGBN064. All constructs were verified by sequencing. Vector pGBN086 for constitutive expression of lmbU-mCherry was prepared by SLICE of linearised vector pGBN054 using primers pG011 and mch_g18A_g21A_Rev and insert amplified on plasmid pGBN062 using primers pG007 and pGBN054_wo_int_RBS_lmrC_U_R.

### Antibiotic susceptibility tests

Minimal inhibition concentration (MIC) values were determined MH agar with a serial twofold dilution of antibiotics. Spores or mycelia from 42 h seed culture or 120 h production culture were resuspened in 100 μl 0,9% NaCl to optical density corresponding to 0.5 – 0.7 McFarland (OD_450_ 0.2 – 0.3) and 5 μl was spotted on MH agar with antibiotic and incubated at 30°C for 5 days (Fig. S7a).

### Analysis of lincomycin and celesticetin production

1 ml of supernatant from 42 h seed culture or 160 h production culture (Fig. S7b) was used for solid-phase extraction as follows: An Oasis HLB 3 cc 60 mg cartridge (hydrophilic-lipophilic balanced sorbent, Waters, USA) was conditioned with 3 ml of methanol and equilibrated with 3 ml of water, and then 1 ml of the supernatant of cultivation broth (for lincomycin extraction pH adjusted to 9.0 with ammonium hydroxide) was loaded. The cartridge was washed with 3 ml of water, and absorbed substances were eluted with 1.5 ml of 80% methanol. The eluent was evaporated to dryness, reconstituted in 150 μl of 50% methanol, and centrifuged at 12,045 g for 5 min at room temperature. The extract was then diluted 10× with methanol: water (1:1 v/v) and analyzed by LC-MS, as described below.

### LC-MS analysis of lincomycin and celesticetin

LC analyses of samples were performed on an Acquity UPLC system equipped with the 2996 DAD detector and LCT premier XE time-of-flight mass spectrometer (Waters, USA). Five microliters of each sample were loaded onto the Acquity UPLC CSH C18 LC column (50 mm × 2.1 mm I.D., particle size 1.7 μm, Waters) kept at 40 °C and eluted with a two -component mobile phase, A and B, A was 1 mM ammonium formate pH 9 (for lincomycin detection, prepared by titration of formic acid 98–100%, Merck, Germany with ammonium hydroxide 28–30%, Sigma-Aldrich, Germany) or A was 0.1% formic acid (for celesticetin detection) and B was acetonitrile (LC-MS grade, Biosolve, Netherlands). The analyses were performed with a linear gradient program (min/%B): 0/5, 1.5/5, 12.5/58 followed by a 1.5min column clean-up (100% B) and 1.5 min equilibration (5% B) at a flow rate of 0.4 ml min^-1^. The DAD detector monitored the column effluent in the range 194-600 nm; the mass spectrometer operated in the “W” mode with its capillary voltage set at +2800 V, cone voltage at +40 V, desolvation gas temperature at 350 °C, ion source block temperature at 120 °C, cone gas flow at 50 l h^-1^, desolvation gas flow at 800 l h^-1^, scan time at 0.15 s, and interscan delay at 0.01s. The data were processed by MassLynx V4.1 (Waters). UV chromatograms monitored at 194 nm were used for lincomycin quantitation based on a five-point linear calibration curve, which was constructed from peak areas corresponding to lincomycin. Calibration solutions were prepared by spiking lincomycin authentic standard at the required concentration into lincomycin-free cultivation broth, extracted and preconcentrated as described above. The quantitation parameters were as follows: concentrations used for the calibration curve were 7.56, 15.125, 31.250, 62.5, and 125 mg l^-1^, the correlation coefficient was r^2^=0.995, and the limit of quantification was 7.56 mg l^-1^ (determined as the lowest point of the calibration curve with precision within 10%). Samples from 42 h of cultivation with lincomycin concentrations below the limit of quantitation were examined using MS detection: extracted ion chromatograms at *m/z* 407.2 were evaluated for the presence of lincomycin. The 160 h samples for celesticetin production were also examined using MS detection: extracted ion chromatograms at *m/z* 528.6 were evaluated for the presence of celesticetin.

### Protein digestion for proteomic analysis

0.1 g of mycelia of 40 h seed culture inoculated from fresh spores (Fig. S7b) was lysed in 0.5 ml 100 mM TEAB containing 2% SDC, 10 mM TCEP, and 40 mM chloroacetamide and boiled at 95°C for 5 min. Protein concentration was determined using a BCA protein assay kit (Thermo), and 20 μg of protein per sample was used for MS sample preparation. Samples were digested with trypsin (at a trypsin/protein ratio of 1/20) at 37°C overnight. After digestion, the samples were acidified with TFA to a final concentration of 1%. SDC was removed by extraction to ethylacetate (65), and peptides were desalted on a Michrom C18 column.

### nLC-MS^2^ Analysis

Nano-reversed-phase columns (EASY-Spray column, 50 cm × 75 μm ID, PepMap C18, 2 μm particles, 100 Å pore size) were used for LC/MS analysis. Mobile phase buffer A was composed of water and 0.1% formic acid. Mobile phase B was composed of acetonitrile and 0.1% formic acid. Samples were loaded onto the trap column (Acclaim PepMap300, C18, 5 μm, 300 Å Wide Pore, 300 μm × 5 mm, 5 Cartridges) for 4 min at 17.5 μl min^-1^ loading buffer composed of water, 2% acetonitrile and 0.1% TFA. Peptides were eluted with a mobile phase B gradient from 4% to 35% B in 60 min. Eluting peptide cations were converted to gas-phase ions by electrospray ionization and analyzed on a Thermo Orbitrap Fusion (Q-OT-qIT, Thermo). Survey scans of peptide precursors from 350 to 1400 m/z were performed at 120 K resolution (at 200 m/z) with a 5 × 10^5^ ion count target. Tandem MS was performed by isolation at 1.5 Th with the quadrupole, HCD fragmentation with a normalized collision energy of 30, and rapid scan MS analysis in the ion trap. The MS2 ion count target was set to 10^4^, and the maximum injection time was 35 ms. Only those precursors with charge states of 2–6 were sampled for MS2. The dynamic exclusion duration was set to 45 s with a 10 ppm tolerance around the selected precursor and its isotopes. Monoisotopic precursor selection was turned on. The instrument was run in top speed mode with 2 s cycles (66).

### Proteomic data analysis and interpretation

All data were analyzed and quantified with the MaxQuant software (version 1.6.1.0) (67). The false discovery rate (FDR) was set to 1% for both proteins and peptides and we specified a minimum peptide length of seven amino acids. The Andromeda search engine was used for the MS/MS spectra search against the *Streptomyces lincolnensis* database (downloaded from NCBI on July 2018). Enzyme specificity was set with the C-terminus as Arg and Lys, also allowing cleavage at proline bonds and a maximum of two missed cleavages. Carbamidomethylation of cysteine was selected as a fixed modification, and N-terminal protein acetylation and methionine oxidation were selected as variable modifications. The “match between runs” feature of MaxQuant was used to transfer identifications to other LC-MS/MS runs based on their masses and retention time (maximum deviation 0.7 min), and this was also used in quantification experiments. Quantifications were performed with label-free algorithms described recently. Obtained normalized data were imported to Perseus 1.6.1.3 software (Max Planck Institute of Biochemistry, Munich)(68). All numeric values corresponding to protein intensity were transformed to a logarithmic scale, and all samples were grouped using categorical annotation. Missing values were then replaced by random numbers drawn from a normal distribution of 1.8 standard deviations downshift and with a width of 0.3 of each sample. Differential analysis of proteins abundance was done using volcano plots (p-values by two-tailed t-test, false discovery rate FDR = 0.05, and fudge factor s0 = 0.1). Pearson correlation distance and Pearson p-values were calculated also by Perseus software using default settings. Heat maps of relative abundance of selected proteins were generated from the matrix of protein intensities without imputation of missing values in Microsoft Excel. Proteomic analysis in 40 h was assessed in 5 biological (4 for ΔC without clindamycin) replicates for each sample/treatment and proteomic analysis of WT in 160 h was performed in three biological replicates.

### Proteomic data accessibility

The mass spectrometry proteomics data have been deposited to the ProteomeXchange Consortium via the PRIDE (69) partner repository with the dataset identifier PXD015009.

### RT-PCR

Mycelia from 16 h seed cultures uninduced or induced by clindamycin (0.5 mg l^-1^) (Fig. S7c) 5 ml the culture were harvested by centrifugation (4000 *g*, 15 min, 4°C) and flash-frozen in liquid nitrogen. For the analysis, samples were defrosted and incubated in 1 ml of RNAprotect cell reagent (Qiagen, 5 min, 25°C). Subsequently, cells were centrifuged (4000 *g,* 15 min, 4°C) and the pellet resuspended in 250 μl TE buffer. The suspension was mixed with glass beads (0.1 mm diameter) in a 2:1 ratio and disrupted using a Fast-Prep (MP Biomedicals) program for 1 × 60 s at a speed of 6 ms^-1^. Immediately after the cell lysis total RNA was isolated using TRI Reagent (T9424-100ml, SIGMA), according to manufacturer protocol. Isolated total RNA, resuspended in 100 μl water, was treated with Turbo DNase (Invitrogen) according to manufacturer protocol, followed by an additional step of total RNA isolation using TRI Reagent. The integrity of RNA was controlled by 2% agarose gel electrophoresis. Purity and concentration of RNA were controlled by NanoDrop.

cDNA was synthesized using SuperScript^™^ III Reverse Transcriptase (Invitrogen) according to manufacturer protocol. 1 μl from reverse transcription reaction mix or total RNA was taken to 20 μl of PCR mix using primers indicated (sequences in Table S2). PCR was done using Taq-Purple DNA Polymerase (T107). Following PCR program was used: 96°C −1 min, and then 30 cycles: 96°C – 10 s, 55°C – 20 s, 72°C – 1 min.

### Quantitative RT-PCR

The 16 h clindamycin-induced and uninduced seed cultures were cultivated and incubated in 1 ml of RNA Protect in the same manner as for RT-PCR Total RNA was extracted with an RNeasy RNA isolation kit (Qiagen). The isolated RNA was treated with DNaseI (0,1 U μl^-1^, 30 min, 37°C) and repurified with an RNeasy RNA isolation kit. RNA quantity and quality were checked with a NanoDrop instrument (DeNovix). RNA quantities were normalized to the lowest concentration of RNA in the samples. Quantities of *lmrC, lmbU, lmbN,* and *16S rRNA* transcripts were measured by one-step qRT-PCR (SuperScript^™^ III Platinum^™^ SYBR^™^ Green One-Step qRT-PCR kit) using the following oligonucleotides (10 μM): lmrCf + lmrCr, lmbUf + lmbUr, lmbNf + lmbNr, and 16SrRNAf + 16SrRNAr. The following real-time PCR program was used: 60°C – 3 min, 95°C – 5 min, and then 40 cycles of 95°C – 10 s, 65°C (*lmbU, lmbN, 16S rRNA*) or 63°C (*lmrC*) – 20 s. Ct values of *lmrC, lmbU*, and *lmbN* transcripts, based on the standard curves, were normalized to Ct values of 16S rRNA. Relative expression was calculated as 2^-ΔCt^.

### Western blotting and immunodetection

Seed cultures (20 ml) inoculated with germinated spores were induced with lincomycin (4 mg l^-1^), clindamycin (0.5 mg l^-1^), pristinamycin IIA (4 mg l^-1^), tiamulin (0.125 mg l^-1^), erythromycin (0.5 mg l^-1^), pristinamycin IA (4 mg l^-1^), chloramphenicol (4 mg l^-1^), carbenicillin (4 mg l^-1^) at 8h and cultivation continued until indicated time points (Fig. S7c). Mycelia were harvested by centrifugation (10 min, 4°C, 4,000 g), washed with buffer 1 (50 mM Tris-HCl, 150 mM NaCl, pH 8.0) and resuspended in sonication buffer (50 mM Tris-HCl, 1x protease inhibitor cocktail (Roche), pH 8.0). Following sonication (3 × 40 s, UP200S Hielscher Ultrasonic GMBH), cell lysates were separated by 8% SDS-PAGE and Western blotted onto a PVDF membrane (semidry transfer, 15 V, 45 min, Trans blot Bio-Rad). The membranes were incubated overnight at 4°C in blocking solution 5% (w/v) Blotting-Grade Blocker (BioRad) in PBS-Tween buffer (1× PBS, 0.05% Tween 20) and incubated a further 1 h with primary antibody (rabbit polyclonal anti-LmrC or rabbit polyclonal anti-mCherry antibody) diluted 1:5,000 in 1% (w/v) nonfat dried milk in PBS-Tween buffer. Membranes were washed for 3 × 15 min in PBS-Tween buffer, incubated for 1 h with horseradish peroxidase-conjugated monoclonal anti-rabbit IgG 1:2,000 in 1% (w/v) Blotting-Grade Blocker in PBS-Tween buffer and washed for 3 × 15 min in PBS-Tween buffer. Antibody complexes were detected using Immobilon Western HRP substrate (Merck) on a ChemiDoc^™^ MP (Bio-Rad).

### Antibodies

The anti-LmrC(2) antibody was generated by GenScript USA Inc. by inoculating a New Zealand Rabbit host strain with a peptide-KLH conjugate containing the LmrC peptide CLQRQAQESAGRAAS. The specificity of LmrC antibody was validated using *S. lincolnensis* WT and ΔC mycelium from 16 h seed culture (Fig. S7C) grown in the absence or the presence of lincomycin (LIN, 4 mg l^-1^) or clindamycin (CLI, 0.5 mg l^-1^) (Fig. S3a). The anti-m-Cherry antibody was purchased from Invitrogen (CAT PA5-34974). A ribosomal protein S7-specific antibody was obtained from Mee-Ngan F. Yap (70).

## Acknowledgments

Research on this project was supported by the Grant Agency of the Czech Republic (15-16225Y and P302-12-P632), Charles University Grant Agency (1767418 to L.V.), and the project BIOCEV – Biotechnology and Biomedicine Center of the Academy of Sciences and Charles University (CZ.1.05/1.1.00/02.0109) from the European Regional Development Fund. Further acknowledgments belong to Karel Harant and Pavel Talacko from Laboratory of Mass Spectrometry, Biocev, Charles University, Faculty of Science, where the proteomic and mass spectrometric analyses were performed; Mee-Ngan F. Yap for the S7 antibody; Mervyn Bibb for M1154 strain; and Hee-Jeon Hong for vectors pIJ10257, pIJ6902 and pMS81.

